# Tunable photoinitiated hydrogel microspheres for direct quantification of cell-generated forces in complex three-dimensional environments

**DOI:** 10.1101/2023.03.31.535168

**Authors:** Antoni Garcia-Herreros, Yi-Ting Yeh, Yunpeng Tu, Adithan Kandasamy, Juan C. del Alamo, Ernesto Criado-Hidalgo

## Abstract

We present a high-throughput method using standard laboratory equipment and microfluidics to produce cellular force microscopy probes with controlled size and elastic modulus. Mechanical forces play crucial roles in cell biology but quantifying these forces in physiologically relevant systems remains challenging due to the complexity of the native cell environment. Polymerized hydrogel microspheres offer great promise for interrogating the mechanics of processes inaccessible to classic force microscopy methods. However, despite significant recent advances, their small size and large surface-to-volume ratio impede the high-yield production of probes with tunable, monodisperse distributions of size and mechanical properties.

To overcome these limitations, we use a flow-focusing microfluidic device to generate large quantities of droplets with highly reproducible, adjustable radii. These droplets contain acrylamide gel precursor and the photoinitiator Lithium phenyl-2,4,6-trimethylbenzoylphosphinate (LAP) as a source of free radicals. LAP provides fine control over microsphere polymerization due to its high molar absorptivity at UV wavelengths and moderate water solubility. The polymerized microspheres can be functionalized with different conjugated extracellular matrix proteins and embedded with fluorescent nanobeads to promote cell attachment and track microsphere deformation.

As proof of concept, we measure the mechanical forces generated by a monolayer of vascular endothelial cells engulfing functionalized microspheres. Individual nanobead motions are tracked in 3D and analyzed to determine the 3D traction forces within seconds and without the need for solving an ill-posed inverse problem. These results reveal that the cell monolayer collectively exerts strong radial compression and subtle lateral distortions on the encapsulated probe.

## 1. Introduction

Mechanical forces play crucial roles in regulating physiological functions, including organ development and tissue homeostasis. Cells constantly sense and respond to mechanical cues from their surroundings by exerting forces. In the past two decades, it has become clear that mechanical forces influence embryonic development^1^, cancer progression^2^, or stem cell differentiation^3^. However, the physical and molecular mechanisms by which these forces, applied non-specifically or to specific adhesion receptors^4^ trigger gene expression and affect cellular function are still not completely understood. In particular, there is a need to interrogate cellular forces *in vivo* or in physiologically relevant *in vitro* models with high similarity to living tissues.

The usage of linearly elastic flat substrates seeded with fluorescent beads transformed the field of cell biomechanics by allowing researchers to measure the 2D^5–7^ and 3D^8–10^ traction forces exerted by cells and the intracellular stresses inside thin, continuous cell monolayers adhering to these substrates^11, 12^. Traction force microscopy and monolayer stress microscopy have become widespread methods in cell biology with relatively well-standardized protocols and analysis techniques. Progress in modeling the non-linear mechanics of fibrillar extracellular matrices has made it possible to reconstruct the forces generated by cells in these complex environments^5, 6^. These approaches commonly rely on constitutive laws, combining theoretical models and empirical data inference, that relate strain and stress in the extracellular environment. However, deriving such constitutive relations remains a significant challenge.

An effective strategy to measure mechanical forces in complex multicellular cultures and living tissues consists of fabricating calibrated microscopic (∼10-100 μm) probes and intercalating them into the experimental region of interest^13^. This approach obviates the problem of obtaining the complex strain-stress relationship of the tissue. One of the first systems to report forces in living tissues used fluorocarbon oil droplets^14, 15^, whose physical properties are well characterized. The anisotropic component of the tissue normal stresses was determined by tracking droplet shape deformations via Young-Laplace’s equation. While droplets do not detect shear stresses or isotropic normal stresses (i.e., pressure) due to their fluid, incompressible nature, they have become widely used due to their easy implementation and non-toxic nature for *in vivo* experiments. Liquid droplets can be directly injected into the tissue but require precise control of the surface chemistry to prevent surrounding tissues from interacting with the highly deformable interface to the point of rupture and posterior internalization of the surfactant or oil.

Hydrogel-based micro-spheres offer an alternative to microdroplets for measuring shear and pressure^16, 17^. These micro-spheres can be fabricated in large quantities by generating a suspension of aqueous droplets in oil, where the aqueous phase contains a gel precursor solution. Vortexing the oil and aqueous phases is a simple and effective method to produce droplet suspensions when obtaining a monodisperse size distribution is not crucial^18, 19^. Besides, flow-focusing microfluidic devices yield monodisperse droplet distributions and are becoming increasingly available^20–23^. Most recent techniques are based on the polymerization of acrylamide, but several studies have utilized other materials, such as alginate or gelatin^24, 25^, whose stiffness is harder to control and/or varies within a relatively narrow range, limiting their usability. Polyethylene glycol (PEG) allows for varying stiffness over a wide range and has desirable biocompatibility ^26^, although its hydrolytic degradability and autooxidation activity might affect cell function and make it less suitable for applications requiring long-term cell culture. Stable polyacrylamide (PAAm) microspheres with adjustable size and stiffness can be fabricated and made biocompatible by functionalization. However, obtaining monodisperse distributions involves complicated microfabrication setups depending on highly specialized equipment^27^ and very careful tuning of the hydrogel polymerization process^28^. Thus, despite significant recent advances^13^, there is still a need for high-throughput platforms to produce hydrogel microspheres of tunable properties and tailored surface chemistry using standard wet laboratory equipment.

This manuscript introduces an approach to generate PAAm micro-spheres with customizable surface chemistry and highly controlled size and Young’s modulus. We use in-house manufactured microfluidic devices to produce droplets of a PAAm precursor solution containing the biocompatible, water-soluble photoinitiator lithium phenyl-2,4,6-trimethylbenzoylphosphinate (LAP). Compared to other commonly used photoinitiators, LAP has a high molar extinction coefficient in response to ultraviolet light (and a moderate one at wavelengths ∼370nm that makes it potentially usable for in situ polymerization in vivo) and has a favorable partition coefficient due to its increased water solubility ^29^, providing precise control over the polymerization process. Our microfluidic device is designed to generate droplets of adjustable size in real time by varying the flow rates between the two liquid phases, without modifying device geometry or restarting the experiment. The precursor microdroplets contain fluorescent nanoparticles to track micro-sphere deformations through three-dimensional coherent point drift particle tracking techniques, making it possible to recover 3D cell-generated stresses *in situ* using straightforward analyses of negligible computational cost. Overall, we demonstrate the high-throughput production of cellular force microscopy probes of highly tunable, reproducible properties using relatively uninvolved processes.

## 2. Results

### 2.1 Generation of gel precursor microdroplets

Emulsification offers an effective approach for the high-throughput generation of gel precursor microdroplets. Traditional emulsification techniques have exploited spatially dependent extensional and shear flows to rupture droplets^18, 21, 30^. These flows can be created easily using standard equipment in a typical wet laboratory (e.g., a vortex mixer). However, this approach typically results in highly polydisperse emulsions with wide distributions of droplet sizes^18^. In this study, we used microfluidics to produce monodisperse droplet suspensions containing acrylamide precursor. We designed a microfluidic device that can be fabricated using standard photolithography methods (Fig 1A). The microfluidic circuit (Fig 1B) has a first stage where flow-focusing emulsification^21, 23, 31^ produces initial droplets and directs them to a collecting channel (Fig 1B, first inset). The microdroplet suspension flows through the collecting channel toward a series of consecutive T junctions (Fig 1B, second inset, and Fig 1 C) that provoke geometrically mediated droplet breakup^32^, considerably reducing droplet size without increasing polydispersity of the daughter droplets. The final droplet size can be easily tuned by adjusting the geometry of the junction, the number of T junctions, and the ratio between the flow rates of the oil and aqueous solutions. Of note, the latter can be tuned while continuously operating the device. A high-frame-rate camera was used to visually inspect the process of droplet breakup and collection (Suppl. Video 1). The resulting microdroplets suspended in oil were collected downstream and polymerized using UV light as described below (see Materials and methods). Afterward, centrifugation was used to isolate the polymerized hydrogel microspheres from the oil phase.

**Figure 1:**
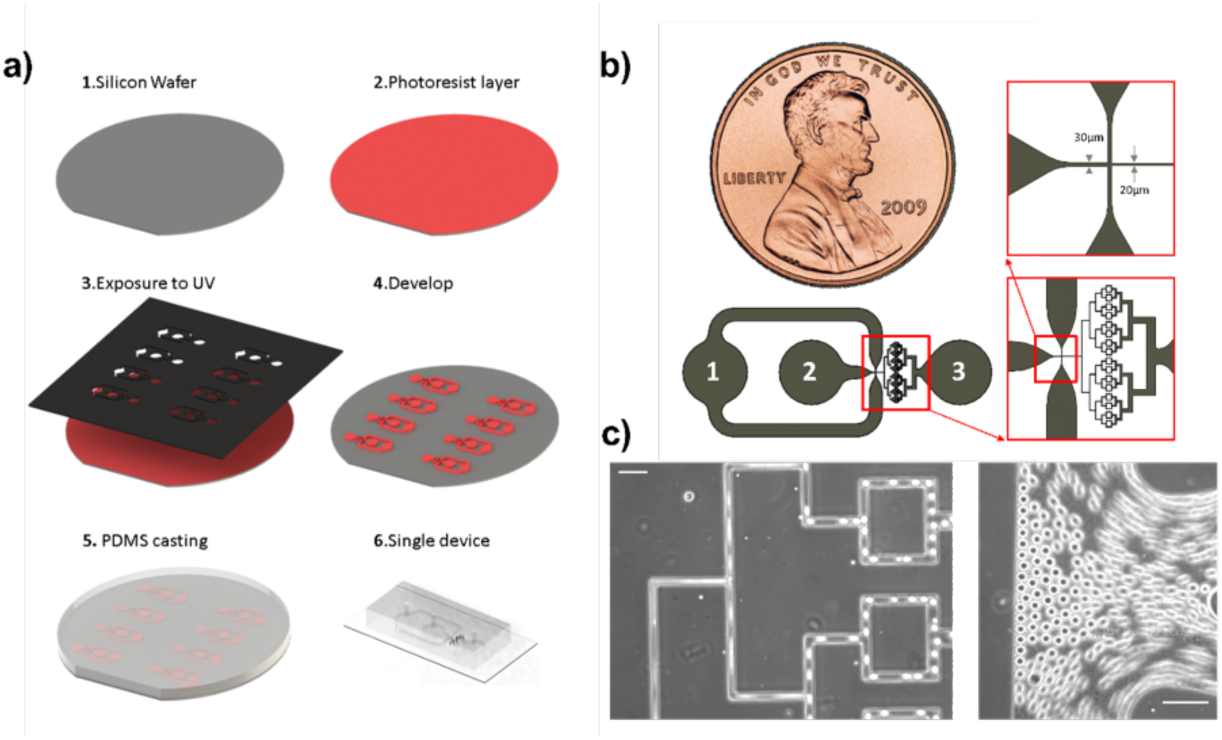
Microfluidic device for high-throughput production of monodisperse gel precursor microspheres via emulsification of water-oil mixtures. **a)** Illustration of the microfluidic device’s multiplexed photolithography fabrication process. **b)** Schematic of the microfluidic device design depicting: 1) Oil phase inlet; 2) Aqueous phase inlet; 3) Outlet of the device from where polyacrylamide beads are collected for further processing. The first (top right) inset shows the flow-focusing stage where the water and oil phases merge and emulsification occurs. The second (bottom right) inset shows a detailed view of the consecutive T-shaped junctions for geometric droplet breakup.

### 2.2 Size distribution of generated microdroplets inside the flow focusing microfluidic device

We characterized the size distribution of the gel precursor microdroplets generated in a microfluidic device with 30-µm-wide channels for the oil and precursor solutions, a 20-µm-wide collecting channel, and flowrates of 60/30 (oil/precursor) µL/h. Droplet size was assessed using their diameter observed using a bright field microscope (Fig 2A). The diameter of the microdroplets produced by the same device was varied in the range 18.6 ± 0.9 – 34.7 ± 1.3 (in μm, mean ± std) by decreasing the oil flowrate from 120 μl/h to 40 μl/h while maintaining the precursor phase flowrate constant at 30 μl/h (Fig 2B). We also generated microdroplets of the same gel precursor aqueous solution in oil using a recently developed process based on inverse emulsification via vortex mixing, which is rapid and does not require microfluidics (Fig 2C)^18^. Vortexing at different rotation speeds allowed for some control over microdroplet size but produced significantly wider diameter distributions (Fig 2D, see also Table 1). For instance, both operating the microfluidic device at 40 μl/h oil and 30 μl/h precursor and vortexing at 3,200 r.p.m. yielded microspheres with a mean diameter of approximately 35 μm; however, the diameter’s standard deviation in the microfluidic device was an order of magnitude lower when compared to vortexing (3.7% of the mean vs. 36%, respectively).

**Figure 2:**
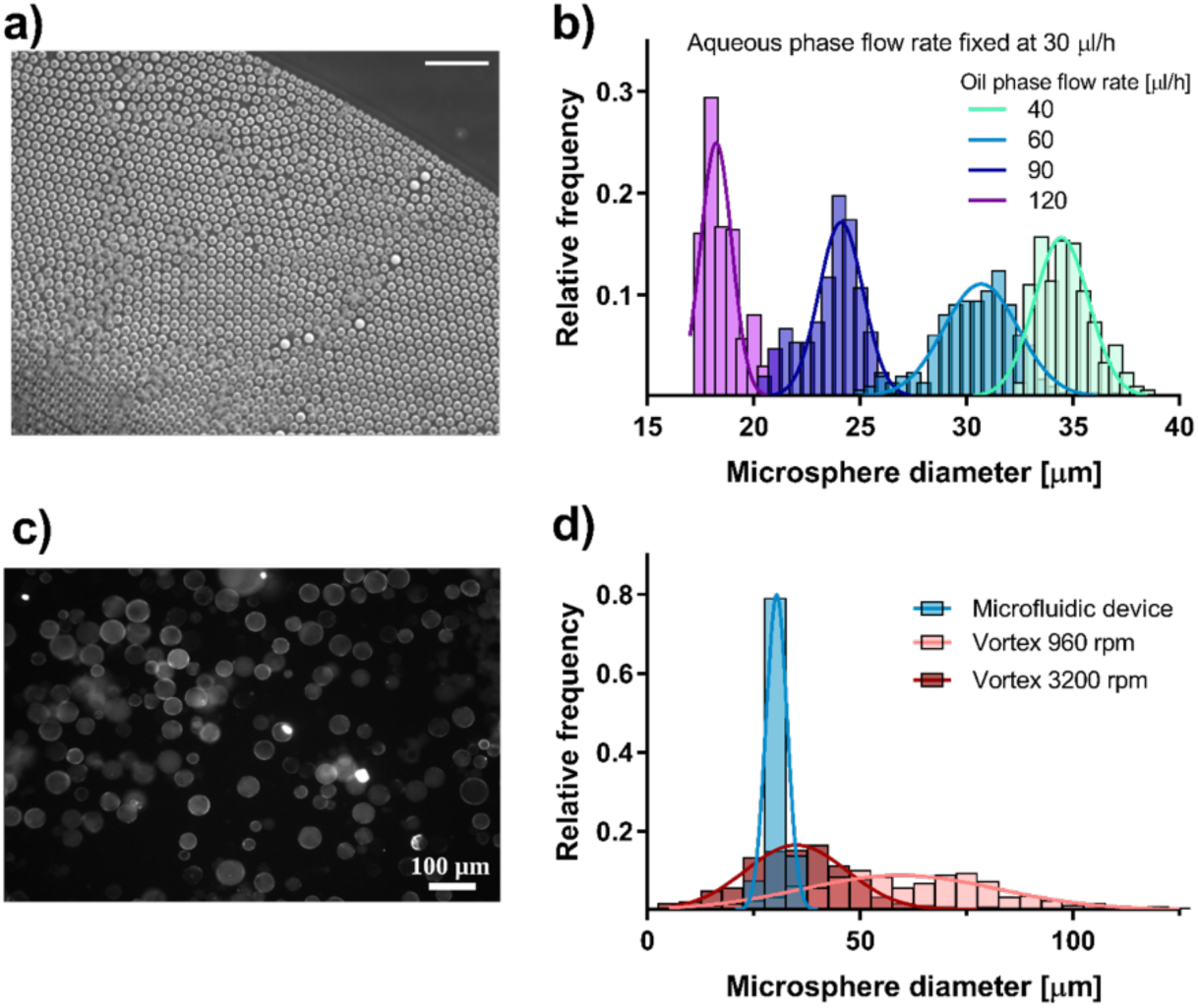
Size distribution of pre-gel droplets produced with a flow focusing microfluidic device. a) Bright field image of droplets produced by the flow-focusing microfluidic device coupled with T junctions shown in Figure 1. Scale bar: 100µm. b) Size (diameter, in microns) distributions of the liquid droplets produced for different oil flow rates and fixed water flow rate. Probability density functions were calculated for at least 100 microspheres in 3 different experiments. c) Microspheres produced by agitating the precursor and oil phases in a vortex mixer for 30 seconds. d) The microfluidic device (blue) produced microspheres with significant lower size variability than the vortex mixer at different speeds (red).

**Table 1:**
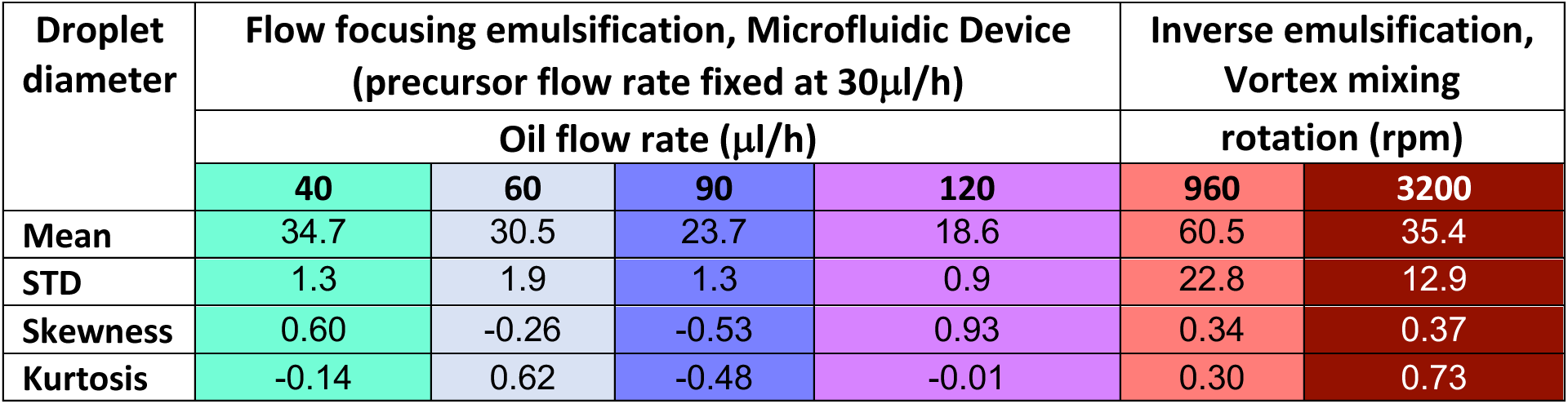
Summarized microdroplet sizes distributions obtained using the microfluidic device and the two-phase mixing technique using the vortex.

### 2.3 Photoinitiated polymerization of precursor acrylamide solution droplets

Once the microdroplets were produced inside the microfluidic device, the presence of surfactant stabilized the emulsion for a significant period, thus preventing droplet coalescence and allowing for droplet polymerization. To control the polymerization, we included the highly efficient, biocompatible, water-soluble photoinitiator lithium phenyl-2,4,6-trimethylbenzoylphosphinate (LAP) in the precursor acrylamide solution. The LAP compound released free radicals when exposed to UV light (302nm) in a benchtop transilluminator, triggering the polymerization reaction inside the droplets. Our rationale for using LAP over other commonly used initiators like ammonium persulfate (APS) or tetramethylethylenediamine (TEMED)^28^ was twofold. First, LAP’s increased water solubility allows a homogeneous polymerization reaction to take place simultaneously all throughout the water-based precursor droplet. Second, LAP’s more favorable partition coefficient enables efficient polymerization of smaller droplets since the concentration of free radicals does not decrease dramatically from the surface of the droplet to its geometric center. Thus, compared with other commonly used substances like ammonium persulfate (APS) or tetramethylethylenediamine (TEMED)^28^, LAP more accurate control of polymerization initiation.

Since carboxylated fluorescent nanobeads were added to the precursor solution to perform 3D traction force microscopy^4, 33^, we used these nanobeads to verify the efficacy and homogeneity of the photoinitiated polymerization process. When pre-gel solution droplets did not polymerize, the fluorescent nanobeads experienced appreciable, uncoordinated Brownian motion (Fig 3A). In contrast, the fluorescent tracers had fixed locations in successfully polymerized microspheres, indicative of a solidified polymeric structure with the nanobeads attached to the polymer backbone (Fig 3B).

**Figure 3:**
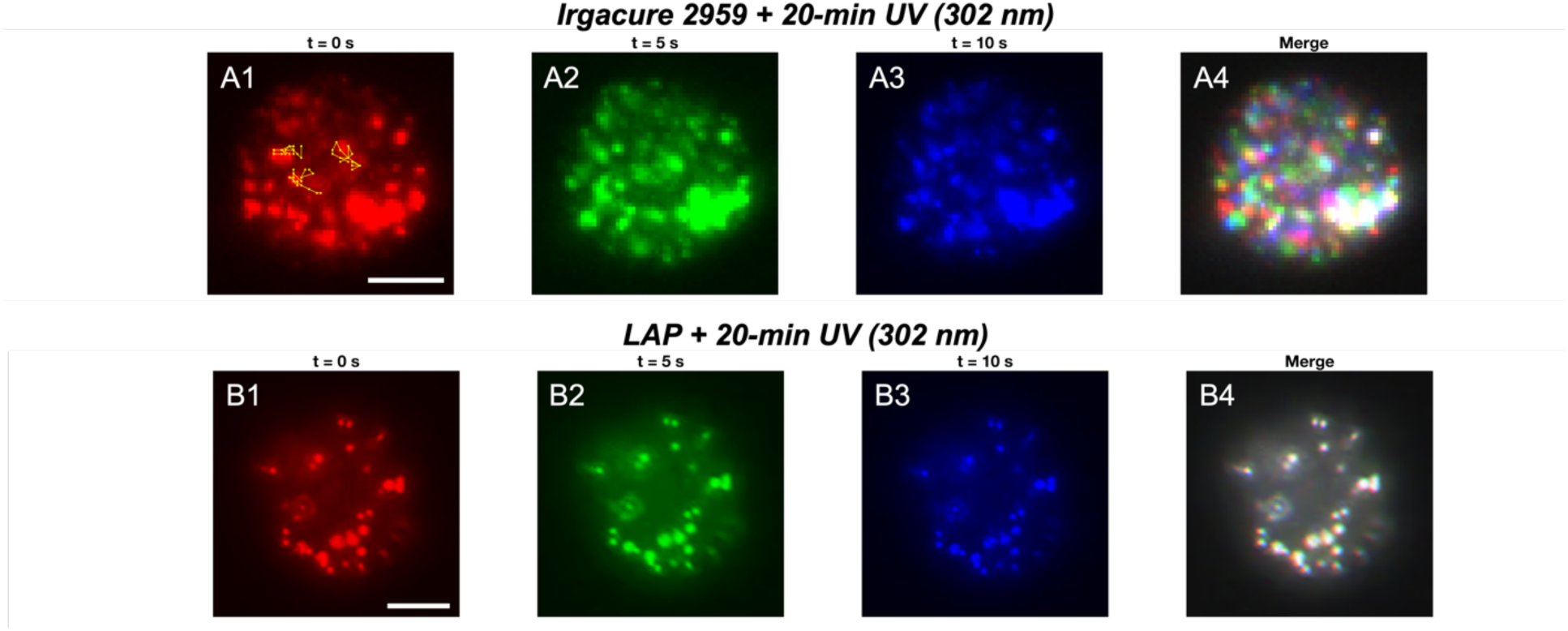
A1-A3) Three snapshots of a microsphere photo-polymerized with Irgacure 2959 after a 20-min exposure to UV light (302nm), taken at 5-second intervals. The Brownian trajectories of three representative nanobeads tracked over 10 seconds are shown in panel A1. A4) Overlaying panels A1-A3 confirm that nanobeads inside the microsphere undergo Brownian motion due to incomplete polymerization. B1-B4) Three snapshots of a microsphere photo-polymerized with LAP after a 20-min exposure to UV light (302nm), taken at 5-second intervals. B4) Overlaying panels B1-B3 indicate complete microsphere polymerization and fixed nanobead positions. Scale bar: 10 microns.

Using this approach, we determined that LAP significantly overperformed Irgacure 2959 (1-[4-(2-hydroxyethoxy)-phenyl]-2-hydroxy-2-methyl-1-propanone), one of the most widely used photoinitiators. While we polymerized microdroplets using concentrations of LAP as low as 0.2 mM, it was practically impossible to polymerize microspheres via UV initiation using Irgacure 2959. Below (see Discussion), we argue this limitation could be due to a combination of factors causing Irgacure 2959 to migrate from the aqueous pre-gel phase to the oil phase before it produces enough free radicals to accelerate the polymerization reaction significantly.

### 2.4 Mechanical characterization of LAP-photoinitiated polyacrylamide gels

The mechanical properties of PAAm gels polymerized using the traditional Ammonium Persulfate (APS) initiator have been described thoroughly^34^. However, the properties of LAP-photoinitiated PAAm gels, such as those used to produce microspheres in our flow-focusing microfluidic devices, have not been characterized. We used atomic force microscopy (AFM) to measure the Young’s Modulus of reference, planar, 12 mm-diameter, 20 μm-thick gel pads (see diagram in Fig 4A) obtained with different ratios of acrylamide, bis-acrylamide, and LAP. For reference, we also measured the Young’s modulus of similar gels using the standard chemical initiator APS. We used three acrylamide/bis-acrylamide ratios known to produce gels of low (*E* ≈ 3 *kPa*), intermediate (*E* ≈ 9 *kPa*), and high (*E* ≈ 40 *kPa*) stiffness (see Materials and Methods Section for details). We carried out these measurements on flat gel pads instead of the microspheres for two reasons. First, it was not possible to manufacture PAAm microspheres of homogeneous stiffness using more standard initiators like APS or Irgacure 2959 to perform the control stiffness measurements. Second, the calculation of Young’s modulus from AFM tip indentation assumes that the gel has an infinitely wide planar shape, producing artifacts in samples with finite lateral size ^35^.

**Figure 4:**
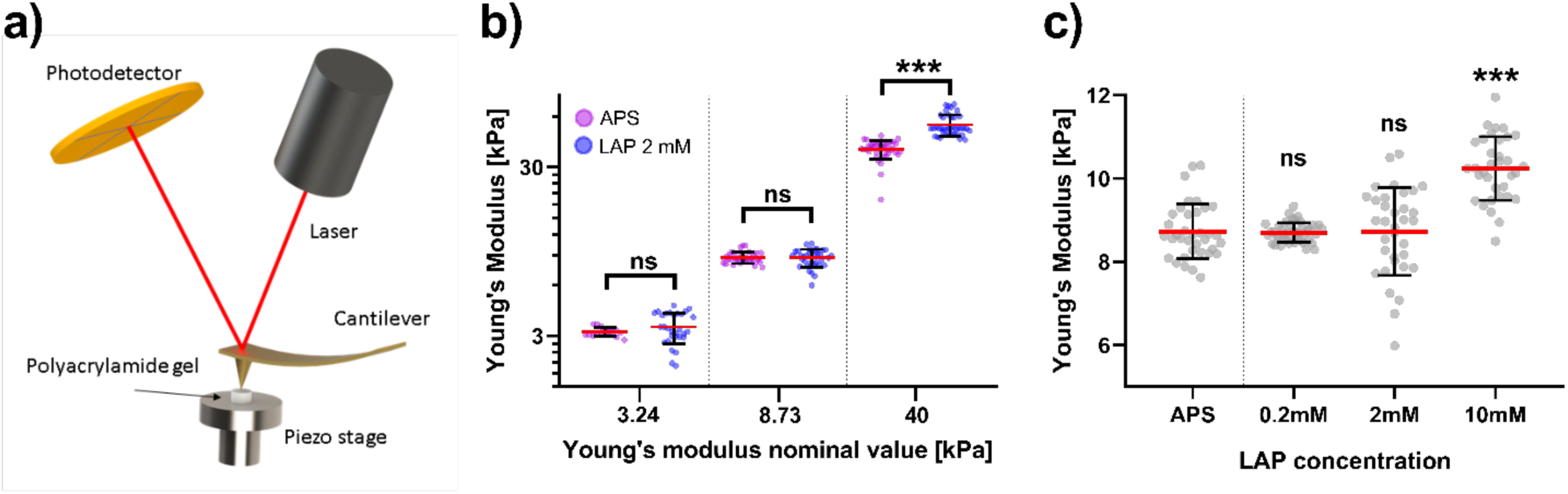
Polyacrylamide gel elasticity characterization by AFM indentation. a) A schematic of the AFM method used to measure the elasticity of the hydrogels. b) Comparison of LAP photoinitiator with a chosen concentration of 2mM with traditional APS 0.1% (w/v) formulation for three different stiffness values. Only for gel stiffness values as high as 40kPa significant differences were observed between the use of APS and LAP. c) Effect of increased concentrations of photoinitiator in the pre-gel solution. Only significant differences were observed when concentration values were increased up to 10 mM. Statistically significant differences were calculated for at least 20 gels in each condition and determined using an unpaired t-test with Welch’s correction (***, p< 0.001).

For gels with low or intermediate stiffness, we obtained statistically indistinguishable values of the Young’s modulus using 2 mM LAP or APS as initiators (Fig 4B). On the other hand, for stiffer gels, there was a small albeit statistically significant difference, with LAP producing stiffer gels than APS (53.4 ± 7.8 vs. 38.3 ± 4.8, p-value <0.001, t-test with Welch correction). In the intermediate stiffness case, which corresponds with a Young’s modulus value customarily used in traction force microscopy^36^, we also evaluated the effect of LAP concentration on gel stiffness keeping constant the ratio of acrylamide to bis-acrylamide concentrations (Fig 4C). Our measurements showed that varying the LAP concentration between 0.2 mM and 2 mM had no statistically significant effect on the stiffness of LAP-initiated gels, which also agreed with the reference, APS-initiated gel stiffness. Nonetheless, when LAP concentration in solution was increased to 10 mM, the Young’s modulus increased when compared with the nominal value. While this increase was moderate, it was statistically significant (10.2 ± 0.7 vs. 8.7 ± 1.1, p-value <0.001, t-test with Welch correction). Overall, these data suggest that there is a wide range of concentrations for which using LAP as photoinitiator does not significantly affect the well-documented mechanical properties of polymerized acrylamide/bis-acrylamide mixtures^34^.

### 2.5 Spatial distribution of fluorescent nanobeads inside PAAm Microspheres

Fluorescent carboxylated nanobeads of ∼ 200 nm in diameter were embedded inside the PAAm microspheres through the precursor solution. Given that these nanobeads serve as fiduciary markers of microsphere deformations, achieving a homogeneous spatial distribution of nanobeads inside the final polymerized microspheres is crucial to ensure high-quality deformation measurements. To characterize this spatial distribution, we imaged the fluorescent nanobeads in our PAAm microspheres and compared their spatial distribution with that obtained in larger (12 mm-diameter, 20 μm-thick) planar PAAm gel pads fabricated for traction force microscopy using well-established methods^33, 36, 37^. Specifically, we took confocal image z-stacks and measured the density of nanobeads per unit area within each imaged x-y slice (e.g., Fig 5A) vs. the position of the slice within the gel.

**Figure 5:**
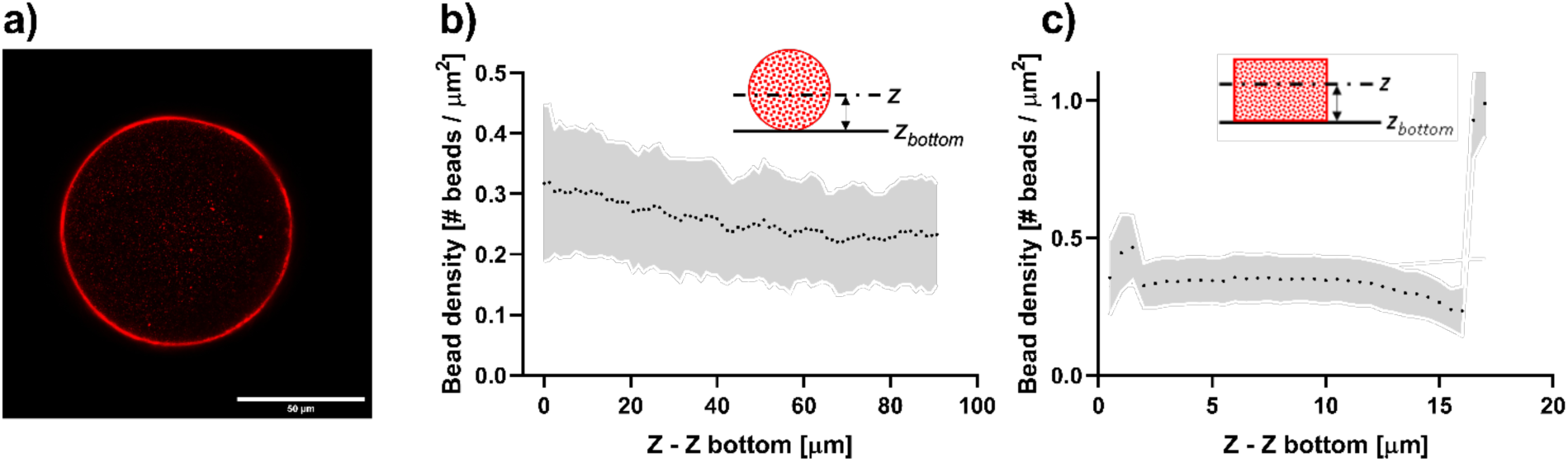
Comparison of the fluorescent microbead distributions in traditionally employed 2D planar PAAm gels vs. spherical photoinitiated polymerized PAAm gels generated with our flow-focusing device. **A)** Image of the equatorial plane of a microsphere. **B)** Density distribution of fluorescent nanobeads for different *Z = constant* planes of the PAAm microsphere. **C)** Density distribution of fluorescent nanobeads for different *Z = constant* planes of a flat PAAm gel slab.

Our measurements showed that the distribution of nanobeads was almost uniform, averaging between 0.32 ± 0.12 µm^-2^ at the bottom of the spheres and 0.23 ± 0.09 µm^-2^ at their top (Fig 5B). This nanobead density compares well with the density we found in large planar gel pads (Fig 5C), with a fairly constant value of 0.3 beads/µm^2^, except near the bottom layer in contact with the coverslip and the top free surface. The bead density at the top free surface increased dramatically to ≈ 1 µm^-2^, a phenomenon that has been observed previously and is often sought after when performing traction force microscopy on flat surfaces^38^. This concentration of nanobeads at the gel’s free surface was also present in the microspheres and is observed, *e.g.*, by the sharp brightness increase at the microsphere’s edge in Figure 5A. However, its effect was averaged out in the density plot in Figure 5B because, in contrast to flat substrates, no single plane slice of the confocal z-stack captured the whole outer surface of the microsphere.

### 2.6 Microsphere surface functionalization and protein conjugation

Polyacrylamide resists the adsorption of proteins and the adhesion of cells. Thus, prior to using PAAm microspheres to study cellular interactions, we treated the microspheres following procedures previously described for planar gel pads^34^ (see Methods Section). Figure 6A-D shows the same microsphere embedded with red (TRITC) fluorescent carboxylated 200-nm-diameter beads and functionalized with 50 µg/mL of Intercellular Adhesion Molecule 1 (ICAM-1), a protein present in the leukocyte extracellular membrane. We co-cultured these microspheres with human umbilical chord vascular endothelial cells (HUVECs) overnight, which were fluorescently stained with green (FITC) phalloidin to label their F-actin. Confocal microscopy images obtained after 12h of co-culture show that endothelial cells formed a confluent monolayer at the bottom substrate (Fig 6E-F) and engulfed the microsphere (Fig 6G-H). These data show a homogeneous distribution of protein coating and confluent cell adhesion to the microsphere surface, demonstrating the efficacy of the protein conjugation protocol. Further SEM imaging of HUVEC-engulfed microspheres (Fig 6I) revealed thick straight filaments, presumably actin-rich stress fibers, distributed radially from the endothelial substrate and towards the center of the microsphere. The presence of these filaments is consistent with strong contractile forces being transmitted to the microsphere.

**Figure 6:**
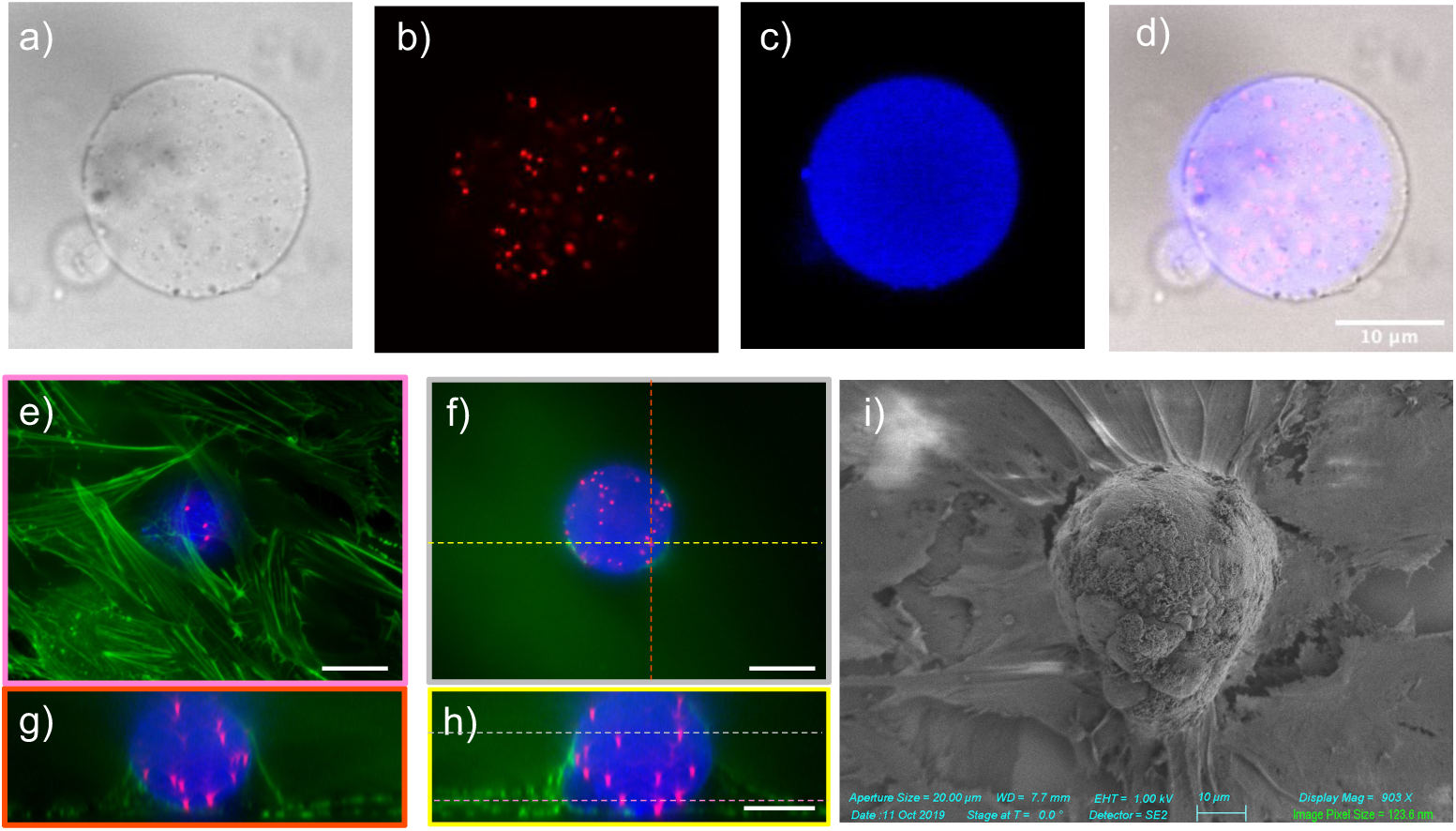
Surface activation and functionalization of polyacrylamide microbeads for cellular interaction. **a)** Bright field of an equatorial plane of a PAAm microsphere. **b)** Red fluorescence image of the same microsphere showing the nanobead distribution. **c)** Blue fluorescence image of the same microsphere showing ICAM-1 functionalization. **d)** Merged image showing the bright-field, red, and blue channels. These microspheres were deposited on fibronectin coated glass substrates and co-cultured with HUVEC cells. F-actin was fluorescently stained with FITC phalloidin (green) and confocal image z-stacks were acquired. Horizontal views **(e-f)** show confluent HUVEC monolayer on the glass substrate. Lateral views **(g-h)** show HUVECs engulfing the microsphere. **i)** SEM image of a PAAm microsphere partially engulfed by HUVECs.

### 2.7 Proof of Concept: Measuring 3D Cellular Forces During Microsphere Encapsulation by Vascular Endothelial Cells

Here, we illustrate the application of hydrogel microspheres to quantify 3D cellular forces by analyzing vascular encapsulation of the microspheres. We generated PAAm microspheres coated with ICAM-1 embedded with fluorescent nanobeads and co-cultured these microspheres with HUVEC cells overnight, as described above. We acquired confocal microscopy images and thentreated the HUVEC cells with Cytochalasin D, a compound that inhibits actin polymerization, to turn the cells into a mechanically relaxed state. We compared the stressed and relaxed states by tracking fluorescent nanoparticle positions to characterize microsphere deformation caused during encapsulation (see Materials and Methods). In short, we ran two instances of Myronenko and Song’s^39^ open-source coherent point drift algorithm for MATLAB. The first run calculated the rigid body motion (translation and rotation) of the nanobeads, which was not considered in the calculation of microsphere stresses. The second run determined the non-rigid motion of the nanobeads after subtracting their rigid body motion. Figure 7A shows segmented nanobeads in the reference relaxed state (magenta) and the deformed state (yellow). To verify the CPD algorithm, Figure 7A also displays the positions of the deformed-state nanobeads after backtracking their CPD-calculated displacement (cyan), showing that they co-localize with the reference-state nanobeads.

**Figure 7:**
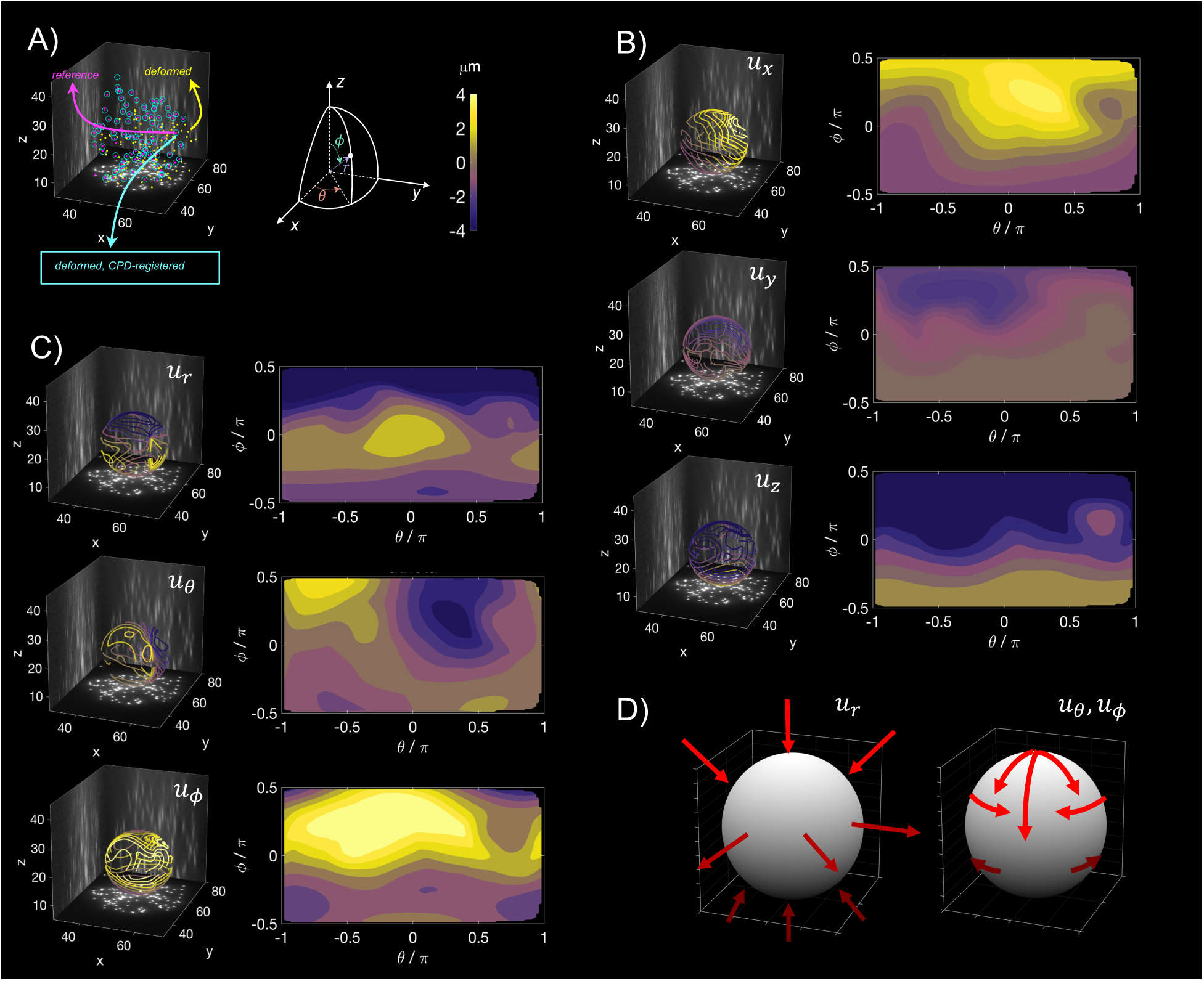
Example of deformation mapping in a microsphere encapsulated by a VEC monolayer. **A) Left:** 0.2-μm-diameter fluorescent nanobeads are segmented from 3D image z-stacks in the deformed (yellow) and undeformed reference (magenta) states. The nanobeads are rendered in 3D together with projections of the cumulative image intensity of the z-stack. The Cartesian (XYZ) axis labels indicate distances in microns. Reversing the measured displacements of the nanobeads in the deformed state yields the cyan circles, which co-localize with the undeformed state nanobeads. **Right:** Schematic illustrating the Cartesian (XYZ) and polar spherical (rθϕ) representations used to map microsphere deformation. **B)** Cartesian nanobead displacements (*u*_*x*_, *u*_*y*_, *u*_*z*_) are interpolated onto a spherical surface of radius R = 10 μm centered at the PAA microsphere’s centroid. Displacement contour levels between - 4 μm and 4 μm are represented in 3D (left) and mapped in 2D vs. azimuth angle θ and elevation angle ϕ (right), and colored according to the color-bar in panel A. **C)** Same as panel C but for displacements projected in polar spherical coordinates (*u*_*r*_, *u*_θ_, *u*_ϕ_). **D)** Schematic summarizing the microsphere deformation pattern.

The nanobead non-rigid motion was then interpolated into a 3D mesh and represented on the surface of the PAAm microsphere, both in Cartesian (Fig 7B) and polar spherical coordinates (Fig 7C). These data indicate that the microsphere was compressed by the encapsulating VECs in its northern polar circle (*u_r_* is large and negative for *φ* ≥ *π*/4 radians or 45°), while it underwent expansion along its equatorial belt (*u_r_* > 0 for −*π*/4 ≲ *φ* ≲ *π*/4) due to Poisson’s effect since PAAm is nearly incompressible. The microsphere’s southern polar circle, in contact with the substrate, also underwent compression, although less significant than in its northern part. In addition to these normal deformations, we measured significant tangential deformations on the sphere’s surface. In the azimuthal direction, the northern hemisphere was characterized by a western *u*_*θ*_ > 0 and eastern *u*_*θ*_ < 0 pattern that created a convergence zone close to the *θ* = 0 meridian, a pattern that was inverted in the southern hemisphere. In the zenithal direction, a relatively intense *u*_*φ*_ > 0 pulled toward the equator in the northern hemisphere while a weaker pattern was observed below the equator. Overall, these tangential and normal deformation patterns are largely consistent with the VEC monolayer adhering to and pulling over the microsphere toward the substrate as summarized in the schematic of Fig 7D. These data illustrate that nanobead-embedded microspheres can resolve cell-generated deformations at the interface between them and the microspheres, including subtle lateral distortions that are not measurable with existing approaches based on oil droplets and hydrogel microspheres with uniformly labeled surface^14, 15, 27^. Nevertheless, it is noted that recording these subtle patterns involves having experimental access to a relaxed reference configuration of the probing microsphere.

A significant advantage of having experimental access to the 3D deformations exerted by the cells on the microspheres is that one can apply Hooke’s law directly to determine cell-generated stresses, since PAAm behaves like a linearly elastic material for a wide range of strain values^40^. In other words, it is not necessary to solve an ill-posed inverse problem to determine the traction stresses. To illustrate this feature, we computed the full 3D stress tensor 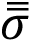 from the deformation measurements of Figure 7, and projected it onto the surface of the microsphere to determine the cell-generated, 3D traction stress vector 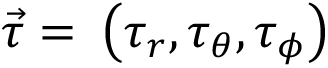 (see Materials and Methods for details). Figure 8 shows the traction stress maps obtained with this procedure. They are consistent with the deformation patterns described above, but, given that the Poisson’s ratio of PAAm is close to incompressible, the normal component *τ_r_* became an order of magnitude higher than the tangential components *τ*_θ_ and *τ*_ϕ_, once again highlighting the ability of resolving subtle lateral distortions with nanobead-embedded PAAm microspheres.

**Figure 8:**
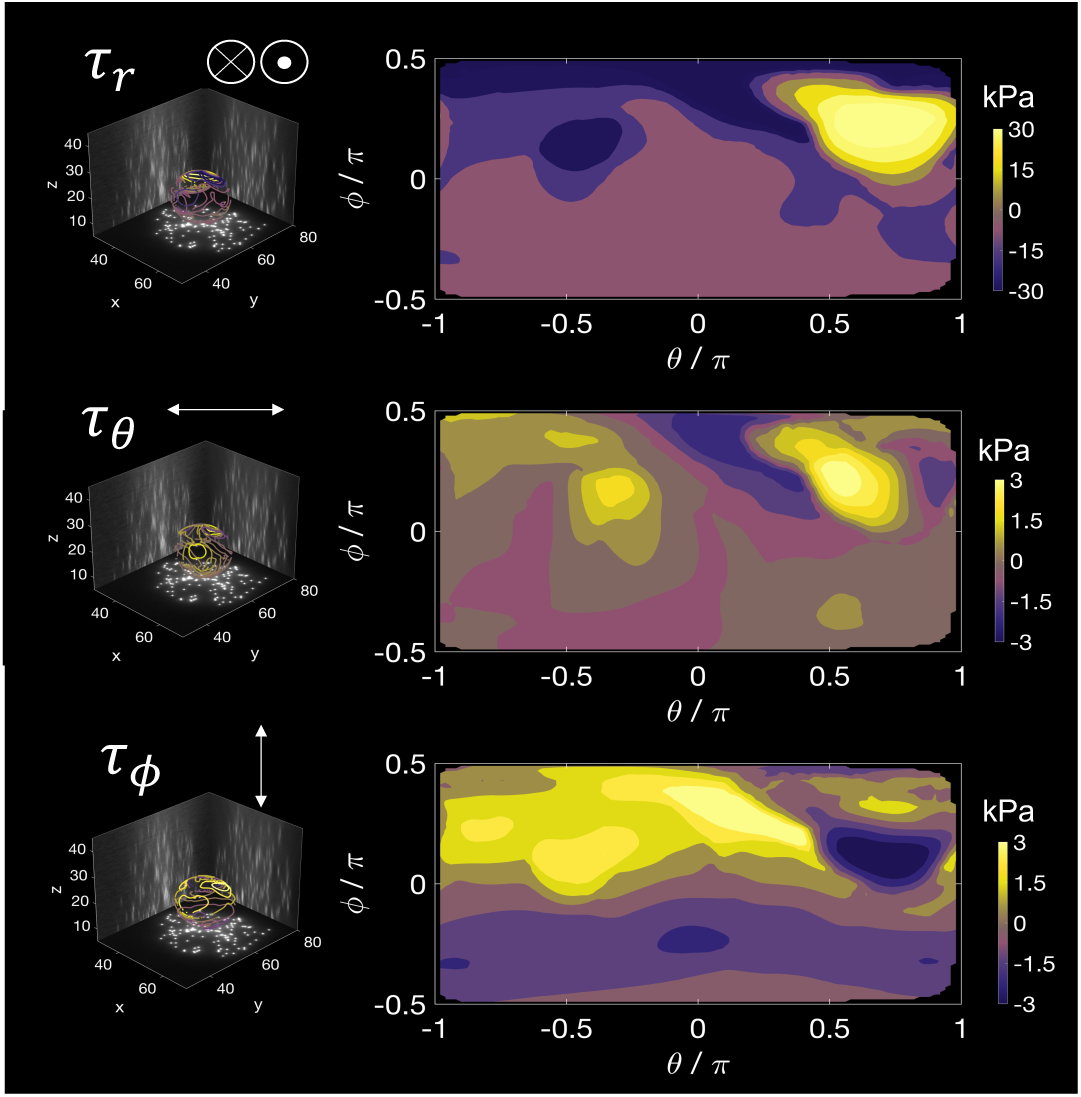
3D traction stresses in a microsphere encapsulated by a VEC monolayer. The 3D deformation field shown in Figure 7 was analyzed using Hooke’s law to compute radial (**top row**), azimuthal (**middle row**) and zenithal (**bottom row**) components of the 3D traction stress vector at the microsphere surface, 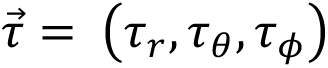 The data are in 3D (left) and mapped in 2D vs. azimuth angle θ and elevation angle ϕ (right), and colored according to the colorbar in panel. Note that the colorbar scale of the *τ_r_* map is different than those of *τ*_θ_ and *τ*_ϕ_.

## 3. Discussion

The *in situ* quantification of mechanical stresses in living tissues and organisms is challenging because of their three-dimensional (3D) geometry and heterogeneous, non-linear, and time-dependent material properties. This makes it difficult to exploit mathematical formalisms to recover stresses from deformation measurements without elaborate modeling assumptions^13, 41^. Concurrent with these issues, it is often impossible to establish the undeformed reference state without disturbing the tissue, *e.g.*, by pharmacological manipulation or laser ablation.

In recent years, microscopic oil and hydrogel probes have been quickly developed to measure mechanical stresses involved in complex biological scenarios spanning a wide range of length scales^14, 15, 24, 27, 28, 42–44^. These probes can be functionalized by surface coating to control the interactions between them and the surrounding cells, and their relaxed state morphology, usually spherical, is determined by their fabrication process. Oil droplet probes can sense anisotropic normal stresses^14^ and, when incorporating a ferrofluid, these probes can also be tuned to perform *in situ* microrheological measurements^15^. However, their incompressibility renders them insensitive to isotropic pressure and their fluid nature limits their suitability to measure shear stresses. On the other hand, hydrogel probes of calibrated elastic properties allow for complete characterization of mechanical stresses^43^. While recent literature anticipates a significant impact for biomechanical stress quantification techniques based on hydrogel probes, implementing these techniques still presents several practical drawbacks that hinder their widespread adoption. In this regard, traction force microscopy of cells seeded on flat polyacrylamide (PAAm) hydrogel substrates, which are relatively straightforward to fabricate and functionalize and provide a precise control of experimental parameters, has set an example by becoming a widely available technique while at the same time increasing in accuracy and resolution over the past two decades^5–10^.

Alginate- or gelatin-based hydrogel microspheres have also been used due to their linearly elastic response and biocompatibility^24, 25^. However, their fabrication process can be relatively complicated, involving several purification steps and careful pH control to ensure the gel’s homogeneity and surface functionalization. Moreover, the stiffness of these hydrogels spans a narrow range in the linear regime, limiting their applicability in biologically relevant scenarios. Other researchers have developed polyacrylamide (PAAm) microspheres, typically generated from emulsifications of a water-oil mixture, to quantify 3D cellular stresses in complex environments^18, 27, 28, 42–44^. In this study, we employed PAAm hydrogels because of the biocompatibility, non-degradability in culture conditions, compressibility, simplicity of surface functionalization, and, most importantly, because the PAAm Young’s modulus can be tuned over a wide range of values by adjusting monomer and crosslinker concentrations^34^.

A handicap common to most existing methods to fabricate PAAm microspheres is the difficulty of controlling the emulsification process to obtain highly reproducible microsphere batches with monodisperse size. Recent efforts have devised an ingenious process of emulsifying acrylamide-bisacrylamide solutions through microporous glass membranes to produce PAAm microspheres with reproducible, tunable dimensions^27^. However, the need for specialized equipment makes this method relatively inaccessible to ordinary laboratories. A more straightforward approach to produce PAAm microbeads relies on inverse emulsification of pre-gel solutions immersed in oil reservoirs^18, 43, 44^, but the need for introducing high levels of mechanical energy during emulsification results in a more widely dispersed distribution of microbead sizes. This work introduces a relatively simple platform to produce PAAm-based microspheres using a flow-focusing microfluidic device. The device combines the flow-focusing strategy to generate the initial droplets and several downstream passive-breakup T junctions that further reduce the size of the initially generated liquid droplets without increasing polidispersity^32^. By varying the oil flow rate, we produced pre-gel droplets of diameters ranging from 20 to 100 µm while keeping the polydispersity index at about 5% at each diameter value.

A second hurdle facing PAAm microsphere fabrication platforms is harnessing the polymerization of the pre-gel droplets to ensure that the final microspheres have the desired mechanical properties. The polymerization of PAAm is customarily triggered by introducing a combination of an initiator and catalyst that accelerates the formation of free radicals in the precursor solution^17^. Initiators such as APS (used in combination with TEMED), commonly used to fabricate flat PAAm substrates for traction force microscopy^34^, have been successfully employed to polymerize microspheres^45^. However, these initiators often generate heterogeneous structural and mechanical properties at the micrometric scale^46^ because they generate free radicals immediately after mixing them with the precursor solution, causing microsphere variability within each production batch^18, 28^.

Photo-activatable initiators like Irgacure 2959 or LAP 3.14, which only generate free radicals when illuminated with wavelengths ≲ 400 nm^29^, allow for better control of PAAm polymerization. Irgacure 2959 is one of the most widely used photo-initiators, but it has a moderate water solubility, reported to be < 2 wt% ^47^, and weak light absorption and free radical production for wavelengths in the near-UV range^29^. In addition, it has a relatively high partition coefficient between the aqueous and oil solvents^48^, *i.e.*, 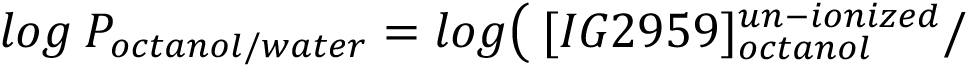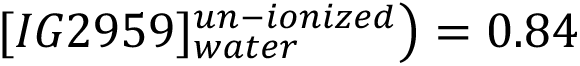, and a relatively low molecular weight (224.3 g/mol) ^49^, implying that it migrates fast from the aqueous phase towards the fluorinated oil phase, when used to polymerize water-based emulsions. Irgacure 2959 has been previously used to control the polymerization of planar, thick (≥ 40 µm) PAAm substrates for cell culture and traction force microscopy^50^. However, these limitations become more critical when trying to polymerize emulsified microdroplets, which have a significantly higher surface-to-volume ratio, rendering PAAm microsphere polymerization via Irgacure 2959 UV initiation challenging in practice.

In our experiments, LAP significantly overperformed Irgacure 2959 in droplet polymerization. Compared to Irgacure 2959, LAP’s higher water solubility, near-UV range absorption, and free radical cleavage made LAP an order of magnitude faster in photopolymerizing hydrogels^29^. Although we could not find reported values for LAP’s partition coefficient in the literature, polymerization of PAAm microspheres was achieved with concentrations of LAP as low as 0.2mM. To validate the mechanical properties of PAAm gels obtained by LAP-mediated photoinitiation, we measured the Young’s modulus of planar substrates using AFM indentation, confirming their linearly elastic behavior and the correspondence between their Young’s modulus and that of PAAm substrates initiated with APS for similar acrylamide/bisacrylamide concentrations. While this similarity held for a wide range of LAP concentrations and PAAm Young’s moduli, stiff gels (*E* ≥ 40 *KPa*) and gels initiated with a high [LAP] ([*LAP*] ≥ 10*mM*) were stiffer than those polymerized with APS, reflecting the higher efficiency of LAP to polymerize PAAm.

One of the many potential applications of hydrogel microspheres could be quantifying the contractile responses of vascular endothelial cells to transmigrating leukocytes^36, 51, 52^, blood flukes and their eggs^53^, or nanoparticles associated with pollution and drug delivery vectors^54^. To investigate these processes in the past, researchers used flat substrates containing fiduciary deformation markers, most often based on micropillar arrays or PAAm gels seeded with fluorescent nanobeads^55, 56^. These experiments revealed significant changes in the traction forces transmitted to the flat substrates while these cells interact with foreign bodies. However, direct measurements of how endothelial cells compress and distort these foreign bodies have been elusive so far. We demonstrated that our microspheres can be functionalized with conjugated ECM proteins (FITC-FN, ICAM-1), which allowed for visualizing their homogenous surface coating and permitted cell attachment and the engulfment of the microspheres by human endothelial cells. We also showed that adding carboxylated fluorescent nanobeads to the precursor solution produces a uniform distribution of embedded nanobeads inside the polymerized microspheres, which can then be used as fiduciary deformation markers. To illustrate the application of nanobead-embedded microspheres as cellular force probes, we determined their full 3D internal deformation fields by tracking the displacements of single nanobeads using an open-source coherent point drift (CPD) algorithm^39^. Knowledge of these fields and the mechanical properties of the microspheres allowed us to determine the complete 3D stress tensor inside the microspheres by applying Hooke’s law directly. In particular, we projected the stress tensor onto the microsphere’s surface to determine the normal and tangential components of the cell-generated traction stresses.

The direct method of determining 3D cell-generated traction stresses circumvents solving an ill-posed inverse continuum mechanics problem, which requires approximating the measurements with theoretical models^27, 57^ or computational analyses (e.g., finite element method)^42^. Also, given its mathematical simplicity, the direct method yields traction stresses within a few seconds, including the computations involved in tracking the nanobead displacements with the CPD algorithm. Compared to previous studies that applied the direct method to alginate microspheres ^24^, using PAAm as probe material ensures a linearly elastic response for a wider range of experimental conditions because of PAAm’s tunable Young’s modulus and more linear stress-strain curve^40^. While it requires recording the nanobead positions at an undeformed reference state, the direct method detects subtle lateral distortions not measurable by reference-free methods that compare the deformed shape of the probe with its ideal, undeformed spherical shape^27^. Even so, the microfabrication process presented in this work can also generate microspheres with fluorescently labelled surfaces for applications where a reference-free approach is critical.

In conclusion, we present a new platform to produce large quantities of photopolymerized hydrogel microspheres with monodisperse sizes, tunable mechanical properties, and custom surface chemistry using a microfluidic device and standard wet laboratory equipment and procedures. By embedding fluorescent nanobeads inside these microspheres, it is possible to measure microsphere strain and recover the 3D cell-generated stresses within seconds following straightforward continuum mechanics concepts and open-source particle tracking software. This technology opens new perspectives for the analysis of stresses exerted not only at the cellular scale, but also inside organoids or developing tissues, where the non-perturbing characterization of mechanical stresses remains challenging to date.

## Materials and Methods

### Design and fabrication of microfluidic emulsification devices

A series of in-house designed and manufactured polydimethylsiloxane (PDMS) microfluidic devices were used to manufacture and collect microdroplets containing the hydrogel precursor solution. These microfluidic devices were manufactured using soft lithography. The master for the microchannel microfluidic device was fabricated in a sequential manner. First, a coating of MCC Primer 80/20 (Microchemicals, Westborough, MA) was applied onto a 4-inch single-side polished silicon wafer (University Wafer, South Boston, MA) and spin-coated at 3000 rpm (accelerating at 1000 rpm/s) over 30 seconds. Shortly after, a first layer of positive photoresist AZ-12XT-20PL-10 (Microchemicals, Westborough, MA) was also spin-coated on top of the primed silicon wafer at 500 rpm over 10 seconds (accelerating at 100 rpm/s). After a short soft bake at 110 °C for 2 min, the sample was exposed to a 375 nm laser (250 mJ/cm^2^) using a MLA150 maskless aligner (Heidelberg Instruments Mikrotechnik GmbH, Germany) fed with the CAD drawings of our device. A post-exposure bake of 1 min at 90 °C was performed, and the features on the wafer were obtained by developing the sample for 2 min using AZ-300MIF (Microchemicals, Westborough, MAs). Two 1-min rinses with deionized water were performed to clear the sample from residues. Profilometry measurements using a Dektak 150 surface profiler (Veeco Instruments Inc. Plainview, NY) confirmed the expected thickness of 20 μm. Subsequently, the wafer was passivated with tridecafluoro-1,1,2,2-tetra-hydrooctyl-1–1trichlorosilane (Gelest, Morrisville, PA) for 15 min inside a vacuum chamber to prevent PDMS adhesion to the wafer in the casting phase of the manufacturing process.

PDMS replicas of the device were made by carefully casting a previously degassed (during 30 min) mixture of the PDMS oligomer and crosslinking agent (Sylgard^®^ 184, Dow Corning Inc. Midland, MI) in a 10:1 (w/w) proportion on top of the passivated silicon wafer. The samples were then cured at 65 °C overnight. The next day the master was peeled off the wafer, cut into several single devices, and the inlet and outlet holes were punched (2.5 mm) with a biopsy puncher (Miltex, Integra Lifesciences, Plainsboro Township, NJ). Finally, to promote PDMS-glass adhesion and sealing, a rectangular coverslip (Corning, 24×60 mm and thickness 1.5 mm) and the PDMS chip were plasma-activated with a UV ozone lamp (Model 30, Jelight Co. Irvine, CA) for 4 min with an oxygen inflow of 0.2 L/min and bonded together at 65 °C for a minimum of 4 hr before they were ready to be used.

### Polyacrylamide microdroplet generation

The hydrogel precursor solution was prepared by mixing 25 µL of 40 % Acrylamide (Millipore-Sigma, Burlington, MA), 30 µL of 2% Bis-Acrylamide (MilliporeSigma, Burlington, MA), 3 µL of 2% 0.2 µm carboxylated FluoSpheres (Invitrogen, Carlsbad, CA), 20 µL of 20 mM Lithium phenyl-2,4,6-trimethylbenzoylphosphinate (LAP, Sigma-Aldrich, St. Louis, MO) and 122 µL of deionized water to a total volume of 200 µL that yielded gels of ∼9kPa. In order to tune hydrogel stiffness the proportion of Acrylamide to Bis-Acrylamide was adequately re-adjusted^34^. This pre-gel solution was kept at 4 °C inside an amber UV-protected 1.5 mL microcentrifuge tube (Eppendorf^®^, Hauppauge, NY), and degassed for 1 h inside a vacuum chamber to reduce oxygen concentration and prevent early gelation inside the driving syringe while running the experiment. The oil phase was prepared by mixing the surfactant oil Krytox^®^ 157 FSH (DuPont, Midland, MI) (10% w), into HFE 7500 Novec Engineered Fluid (3M, Maplewood, MN). Afterward, both the pre-gel and oil solutions were added to different 1 mL syringes and connected to their respective inlets in the microfluidic device using PTFE tubing (1/32”ID and 3/32”OD). The flow rates through the device channels were controlled using two syringe pumps (NE-300 and NE-4000, New Era Syringe Pump). These flow rates were adjusted according to device dimensions, desired droplet size, and droplet generation frequency. For a typical droplet size of ∼35 µm, we used a channel width of 30 µm and flow rates of 60/20 (Oil/Pre-gel) µL/h. The steady state was typically achieved after 15 min, and the device was left running for a minimum of 3 h while droplets were being collected in a 1.5 mL microcentrifuge tube (Eppendorf^®^, Hauppauge, NY).

### Hydrogel precursor microdroplet collection and polymerization

After running the microfluidic emulsification device for 3 h to generate a sufficient amount of microdroplets, the latter were collected directly from the outlet of the microfluidic device inside an amber UV-protected 1.5 mL microcentrifuge tube (Eppendorf^®^, Hauppauge, NY). The solution containing the droplets was transferred into a regular transparent microcentrifuge tube which was exposed to UV (302 nm) light using a benchtop transilluminator (UVP, Analytik Jena US LLC, Upland CA) for 15 min to polymerize the polyacrylamide gel. Subsequently, the oil phase was carefully removed with a pipette tip and replaced with 1 mL of 20% v/v of PFO (1H,1H,2H,2H-perfluoro-1-octanol 97 %, Sigma-Aldrich) in Novec 7500 oil for 3 hr to remove the surfactant from the microspheres. Finally, the solution was centrifuged at 3500 rpm (Sorvall Legend RT, Beckman Coulter Inc. Indianapolis, IN), the oil phase was carefully removed, and the hydrogel microspheres were resuspended in PBS and stored at 4 °C. The centrifugation-resuspension step was repeated three times to achieve improved purity of the resulting microsphere solution in PBS.

### Surface activation and functionalization of polymerized microspheres

Surface activation of hydrogel microspheres for protein conjugation was carried out following the standard procedure used in planar polyacrylamide gel substrates^34^. First, the microspheres were pelleted by centrifugation at 3500 rpm, and then 1 mL of 0.2 mg/ml Sulfo SANPAH (Thermo Fisher Scientific, Waltham, MA), a heterobifunctional protein cross-linker used to covalently bind proteins to polyacrylamide substrates, was added to the microcentrifuge tube. We exposed the sample to UV light (254nm) for 7 min followed by a centrifugation step and posterior wash with PBS 1X. This process was repeated twice. The N-hydroxysuccinimide ester in sulfo-SANPAH can then react with the primary amines of proteins to complete the attachment of proteins to the surface of the gel. To bind the ECM (FN-FITC, ICAM-1) with the activated polyacrylamide microspheres, we incubated the latter in a solution containing 50 µg/mL of the protein conjugate in PBS at 37 °C for a minimum of 3 h.

### Fabrication of polyacrylamide microspheres by inverse emulsification via vortex mixing

Inverse emulsification of the PAAm pre-gel solution was performed by dissolving 50 μL of the solution in 500 μL of HFE 7500 Novec Engineer Fluid (3M, Maplewood, MN) containing (10% w) of the surfactant Krytox® 157 FSH (DuPont, Midland, MI) inside a 1.5 mL Eppendorf tube. Subsequently, the mixture was degassed for 15 min inside a vacuum chamber, then vigorously stirred for 1 min using a vortexer at maximum speed^18^. Immediately after that step, the Eppendorf tube was directly exposed to UV (302 nm) light with a benchtop transilluminator (UVP) for 15 min to initiate polymerization. Once polymerization was completed, the supernatant oil phase was carefully removed with a pipette tip, and the remaining solution containing the hydrogel microspheres was supplemented with 1 mL of 20% v/v of PFO (1H,1H,2H,2H-perfluoro-1-octanol 97 %, Sigma-Aldrich) in Novec 7500 for 3 h to remove the surfactant from the droplets. Right after, the solution was centrifuged at 3500 rpm (Sorvall Legend RT, Beckman Coulter Inc. Indianapolis, IN), the supernatant oil phase was carefully removed, and the hydrogel microspheres were resuspended in PBS and stored at 4 °C. The centrifugation-resuspension step was repeated three times to improve the purity of the resulting microsphere solution in PBS. The subsequent surface activation and functionalization were performed in the same manner as described above.

### Flat Polyacrylamide Substrate Fabrication

Collagen-coated PAAm gel pads of size 12 mm x 20 μm (diameter x thickness) were prepared as previously described by our group for traction force microscopy^36, 58^. Basically, square 25-mm coverslips were activated inside a UV ozone lamp (Model 30, Jelight Co. Irvine, CA) for 10 min and then coated with a 1M solution of NaOH in deionized (DI) water for 5 min. After this step, the coverslips were washed twice with DI water and completely dried it using a vacuum line. The surface of the coverslips were then coated with a solution of 3% (v/v) (3-Aminopropyl) triethoxysilane (APTES, Sigma-Aldrich, St. Louis, MO) in ethanol for 20 min at room temperature. After this step, the coverslips were rinsed with pure ethanol and soft-baked for 5 min at 37 °C. Once the coverslips were ready, the pre-gel solution was prepared with a mixture of 5% Acrylamide and 0.3% Bisacrylamide (calibrated elsewhere^34^ to yield a Young’s modulus E = 8.7 kPa), and seeded with 0.03% carboxylated 0.2-μm diameter FluoSpheres (Invitrogen, Carlsbad, CA). The solution was then mixed with 1% (v/v) of 10% (w/v) Ammonium Persulfate (APS, Sigma-Aldrich, St. Louis, MO) in DI water and N,N,N’,N’-Tetramethyl ethylenediamine (TEMED, Sigma-Aldrich, St. Louis, MO) to initiate the polymerization reaction. Subsequently, 3 µL of the pre-gel solution was quickly pipetted on top of the silanized coverslips, covered with a round 12-mm coverslip on top, and immediately inverted. The gel was let to polymerize for 30 min. During polymerization, carboxylated fluorescent nanospheres tend to migrate preferentially to the top and the bottom (i.e., the free surfaces of the gel). After polymerization, a 0.15 mg/mL Sulfo-SANPAH (Thermo Fisher Scientific, Waltham, MA) solution in DI water was used to cover the gels, followed by UV activation to facilitate the cross-linking of 125 μg/mL of rat tail Collagen Type I (Dow Corning Inc. Midland, MI) to the surface of the polyacrylamide. The gels were incubated for 1 h at 37 °C and then equilibrated with medium for at least 3 h. We measured the thickness of the substrates by locating the top and bottom planes of the gel and subtracting their vertical positions as previously described^36^. The Poisson’s ratio of the gel was measured to be *ν* = 0.46, following an elastographic traction force microscopy method previously developed by our group^59^.

### Imaging

Three-dimensional z-stacks of bright-field and fluorescent images were taken using an Olympus IX81 confocal microscope (Olympus Corp. Tokyo, Japan) with a cooled CCD camera (Hamamatsu Photonics, Shizuoka, Japan), using Metamorph software (Molecular Devices LLC. San Jose, CA) and a 40X, 1.35-N.A. oil-immersion objective. Higher resolution imaging for single microspheres study was performed on an enclosed Zeiss LSM 880 confocal microscope (Carl Zeiss AG, Oberkochen, Germany) with superresolution capabilities under the fast Airyscan mode. A 40x water lens was used. For experiments solely related to characterizing PAAm microsphere shapes, a 200-µm thick z-stacks were acquired with either 1.21 µm or 2.21 µm spacing between images in the z direction. The in-plane calibration factors were 0.0824 µm/px and 0.123 µm/px, respectively. Up to 4 different fluorescent channels were acquired: DAPI, FITC, TRITC and CY5.

In addition, we took scanning electron microscopy (SEM) images of select samples. To prevent the presence of organic contaminants inside SEM instrument, we fixed the samples using a fixative buffer composed of 4% Paraformaldehyde (PFA, Biotium, Freemont, CA) supplemented with 2.5% of Glutaraldehyde (Sigma-Aldrich, St. Louis, MO) in Phosphate Buffered Saline (PBS). We first aspirated the medium in our samples, covered them completely with the fixative buffer and incubated overnight at 4 °C. We replaced the buffer solution with a sequence of increased concentration ethanol solutions (50, 60, 75, 85, 90 and 100%) and left them soak for 15-20 min. The samples were then dehydrated and ready for drying. To prevent collapse of biological samples due to surface tension, Critical Point Drying (CPD) was performed on all samples following dehydration using a Tousimis AutoSamdri 815A critical point dryer (Tousimis, Rockville, MD). After CPD, the samples were ready for sputter coating, which was performed on a Emitech K575X Iridium Sputter Coater. Imaging was then performed using a FEI Quanta FEG 250 SEM with environmental capabilities. Images were acquired at 1kV and with magnifications ranging from 441X to 16,770X.

### Atomic Force Microscopy Measurement of PAAm Hydrogel Young’s Modulus

Hydrogels of varying stiffness were prepared by mixing 40% Acrylamide (Millipore-Sigma, Burlington, MA), 2% Bis-Acrylamide (MilliporeSigma, Burlington, MA), 20 mM stock solutions of Lithium phenyl-2,4,6-trimethylbenzoylphosphinate (LAP, Sigma-Aldrich, St. Louis, MO) and water in the appropriate concentration. Three different concentrations were chosen a priori based on a previous calibration of Young’s modulus vs. concentration in PAAm gels polymerized using ammonium persulfate (APS) and Tetramethylethylenediamine (TEMED)^34^. Flat gel pads were prepared by filling a cylindrical cavity formed by two previously treated coverslips (20 x 20 mm, Sigma Aldrich) and a circular stainless-steel washer. This setup allowed us to produce 5-mm thick gels to prevent possible interference with the bottom substrate when performing nanoindentations^9^. All samples were polymerized by exposing them to UV (302nm) with a benchtop transilluminator (UVP, Analytik Jena US LLC, Upland CA) for 1 minute. After this step, the hydrogel pads were soaked in deionized water and kept at 4°C to prevent evaporation. All the measurements were carried out during the following 48 h to prevent the aging of the hydrogels.

The elasticity of the PAAm hydrogels was characterized using a Scanning Probe Microscope (Veeco Instruments Inc. Plainview, NY) with a Nanoscope IV controller. Nanoindentation curves were obtained using a standard pyramidal tip cantilever (k = 0.24N/m, DNP-10, Bruker Nano Surfaces, Goleta, CA). The Sader method was employed to refine the manufacturer’s specification of the tip cantilever’s spring constant^60^. To reduce the effect of adhesion forces and prevent evaporation, all experiments were conducted using a fluid cell, where the tip and the sample were covered by deionized water at room temperature. A topographical image of 5 x 5 μm was captured for each gel and a minimum of 15 nanoindentations were performed. Nanoindentation curves were obtained by specifying the total movement of the piezo stage (750 nm) and a maximum deflection threshold relative to the baseline of 35 nm. Forward and reverse velocities were fixed at 50 nm/s, and the scan rate was set to 0.1 Hz. After each sample was characterized, the deflection sensitivity of the cantilever was corrected by performing a nanoindentation curve on a silicon surface where no indentation was possible, and consequently, piezo movement of the stage and cantilever deflection must be equal. The Young’s modulus for the different samples was obtained using the NanoScope Analysis software (Version 1.5, Bruker Nano Surfaces, Goleta, CA). The tip half angle was set to 20° as specified by the manufacturer. A linearized Sneddon model^61^ and its correction for pyramidal tips^62^ were used to fit our data.

### Microsphere deformation and stress quantification

We tracked the motion of the fluorescent nanoparticles embedded into the PAAm microspheres by post-processing confocal image z-stacks taken before (stressed state) and after (reference relaxed state) treating cells with 2 mM Cytochalasin D to disrupt actin polymerization. Image post-processing was performed using custom-written MATLAB scripts. First, we segmented the nanobeads in the image stacks by applying an intensity threshold operation that identified the 1% brightest pixels in the reference stack. We then grouped the spatially connected pixels in the binary image resulting from the thresholding operation and cleaned out objects smaller than 10 pixels to reduce noise. The number of segmented nanobeads was counted, and then, the same operation was performed on the stress state z-stack, this time starting with a threshold 10X higher than in the reference stack, which was iteratively lowered until the number of segmented nanobeads in the stress z-stack matched the number of segmented nanobeads in the reference z-stack. We then identified the 3D centroid positions of all segmented nanobeads in each z-stack, created point clouds from these positions (i.e., using the pointCloud MATLAB function), and filtered out these point clouds to remove outliers (i.e., using the pcdenoise MATLAB function). Subsequently, we ran two instances of Myronenko and Song’s^39^ open-source coherent point drift algorithm (CPD), which was downloaded and compiled to run in MATLAB. The first CPD run was performed using the “rigid” option to determine the collective translation and rotation motion of the nanobeads, which accounts for rigid body motion of the microspheres that does not generate elastic stresses. This run typically takes < 5 s on an ordinary laptop computer for the usual number of nanobeads, *N ∼ 100-200* segmented in a *R ∼ 10-μm* microsphere. The second run was carried out with the “nonrigid” option to determine nanobead motion that accounts for elastic microsphere deformations. This second run typically takes < 10 s on an ordinary laptop computer. The results of the CPD algorithm were mostly insensitive to variations in the operating parameters of the algorithm within reasonably wide ranges centered at the values provided in Myronenko’s software suite examples.

The resulting nanobead non-rigid motions accounting for microsphere deformation were interpolated into a uniform 3D mesh (i.e., using the griddata MATLAB function). Interpolated deformation fields were differentiated using 2^nd^-order centered finite differences, and the resulting strains, 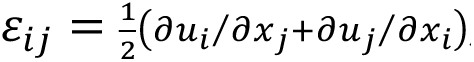, were used to determine the 3D stress tensor inside the microsphere by applying Hooke’s law

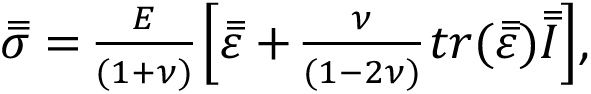

where 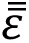 is the strain tensor, 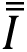 is the second-rank identity tensor (i.e., the 3 x 3 identity matrix), and *E* = 9*kPa* and *ν* = 0.46 are respectively the Young’s modulus and Poisson’s ratio of the PAAm gel. The interpolation and differentiation of the measured deformations were also performed in polar spherical coordinates (*r*, *θ*, *φ*) to facilitate expressing the deformations in the directions normal (*r*) and tangential to the microsphere’s surface (*θ*, *φ*), thus allowing to independently evaluate the expansion/compression and the distortion of the microsphere caused by the cells. Likewise, we projected the 3D stress tensor onto the surface of the microsphere to determine the 3D traction stress vector 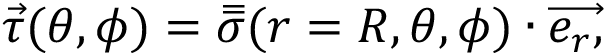 where 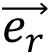 is the unit vector in the radial direction and expressed this traction stress tensor in polar coordinates.

## Notes

### Competing Interest Statement

The authors have declared no competing interest.

